# How Interactions During Viral-Viral Coinfection Can Shape Infection Kinetics

**DOI:** 10.1101/2023.04.05.535744

**Authors:** Lubna Pinky, Joseph R DeAguero, Christopher H Remien, Amber M. Smith

## Abstract

Respiratory virus infections are a leading cause of disease worldwide with multiple viruses detected in 20-30% of cases and several viruses simultaneously circulating. Some infections with viral copathogens have been shown to result in reduced pathogenicity while other virus pairings can worsen disease. The mechanisms driving these dichotomous outcomes are likely variable and have only begun to be examined in the laboratory and clinic. To better understand viral-viral coinfections and predict potential mechanisms that result in distinct disease outcomes, we first systematically fit mathematical models to viral load data from ferrets infected with respiratory syncytial virus (RSV) followed by influenza A virus (IAV) after 3 days. The results suggested that IAV reduced the rate of RSV production while RSV reduced the rate of IAV infected cell clearance. We then explored the realm of possible dynamics for scenarios not examined experimentally, including different infection order, coinfection timing, interaction mechanisms, and viral pairings. IAV coinfection with rhinovirus (RV) or SARS-CoV-2 (CoV2) was examined by using human viral load data from single infections together with murine weight loss data from IAV-RV, RV-IAV, and IAV-CoV2 coinfections to guide the interpretation of the model results. Similar to the results with RSV-IAV coinfection, this analysis showed that the increased disease severity observed during murine IAV-RV or IAV-CoV2 coinfection was likely due to slower clearance of IAV infected cells by the other viruses. On the contrary, the improved outcome when IAV followed RV could be replicated when the rate of RV infected cell clearance was reduced by IAV. Simulating viral-viral coinfections in this way provides new insights about how viral-viral interactions can regulate disease severity during coinfection and yields testable hypotheses ripe for experimental evaluation.

## Introduction

During a respiratory infection, multiple viruses may be present and working in concert to cause disease [1–5]. Several respiratory viruses, including rhinovirus (RV), respiratory syncytial virus (RSV), influenza A and B viruses (IAV and IBV), human metapneumovirus (HMPV), human parainfluenza viruses (PIV), adenoviruses (ADV), and coronaviruses (CoVs), have been found concurrently within hosts with pneumonia [6–20]. Children are more likely to be infected with multiple respiratory viruses concurrently, where up to 30% have more than one respiratory virus present when admitted to the hospital for severe clinical disease [16, 19, 21–25]. Patients with coinfection have shown diverse disease outcomes that ranged from mild to severe with severity increasing compared to patients who were infected with a single virus.

Data on viral-viral coinfections is somewhat limited, but some studies have begun evaluating outcomes of coinfection with commonly observed viral pairs (e.g., IAV and RSV [26–31], IAV and PIV [32], IAV and RV [28,33–35], RSV and RV [28], RSV and hMPV [36], IAV and severe acute respiratory syndrome (SARS)-CoV-2 [37–43], RSV and SARS-CoV-2 [43], and RV and SARS-CoV-2 [44]). Collectively, these studies observed diverse outcomes of viral-viral coinfections with some interactions resulting in enhanced spread of one or both viruses within the respiratory tract while other mechanisms seem to work in an inhibitory manner. For example, PIV-2 was shown to enhance cell-to-cell fusion through expression of its surface glycoproteins, which boosted viral spread between cells and increased IAV titers but not PIV titers [32]. On the contrary, IAV was shown to limit a concurrent RSV infection by promoting intracellular competition for proteins or amino acids needed for the successful replication of both viruses within cell cultures [11, 26]. *In vivo* infections in animal models support the exclusion of RSV by IAV, and suggest that RSV prior to IAV decreases disease severity [27, 29, 30]. A similar competitive exclusion was observed in 3D tissue cultures during coinfection with RSV and hMPV, where hMPV was inhibited without any effect on RSV [36]. The reduction was accompanied by higher type I and III interferon (IFN) responses [36]. IFN-mediated effects were also implicated in RV-IAV coinfection where RV-induced IFN protected against subsequent IAV infection within differentiated airway cell cultures [34].

The relative timing between viruses and the order in which the viruses infect the host seem to contribute to differing disease outcomes [27, 29, 33, 37, 39, 40, 42]. Interestingly, the exclusion effect during RSV-HMPV coinfection was more robust during a concurrent infection compared to an infection where HMPV followed RSV after two days. This is in contrast to other viral-viral coinfections where infections separated by 2 to 5 d had more robust effects [33, 37, 42]. For example, RV was shown to attenuate IAV-mediated disease severity and reduce IAV titers when RV infection occurred 2 d before IAV, but the effect was reduced during simultaneous infection [33]. Conversely, animals coinfected with IAV 2 days before RV experienced greater disease severity [33]. Similar outcomes occurred in animals coinfected with IAV 3 d before SARS-CoV-2 [37].

These empirical studies illuminate the breadth of interactions that lead to diverse outcomes of viral-viral coinfections and the need for mathematical methods that can dissect complex, time-dependent, and potentially nonlinear mechanisms. Viral dynamics models have contributed significantly to the understanding of biological mechanisms underlying respiratory virus infections. Our study examining the resulting dynamics from two viruses competing for target cells [45] suggested that a faster replicating virus would outcompete other viruses. However, it is possible that viruses infect different cells or infect different areas of the respiratory tract. In addition, as noted above, they may also inhibit or enhance other processes (e.g., replication rates and/or immune responses) and, ultimately, modulate disease. Unfortunately, current empirical studies on viral-viral coinfections often lack quantitative information on viral loads and/or host immune responses. This creates a challenge to define mechanisms and for use of mathematical modeling approaches, which typically include fitting a mechanistic model to data. However, in most murine studies, weight loss, which is a measure of disease severity, is tracked, and our recent work showed that mathematical models can accurately connect animal weight loss to infection kinetics and lung inflammation in single infections and bacterial coinfections [46, 47]. These links allow us to better interpret weight loss data and afford modeling studies that ability to assess mechanisms with limited data.

To better understand the underlying mechanisms and determine how different virus orders, timings, and pairings affect the infection dynamics and disease severity, we first paired two mathematical models, which have differing hypotheses about whether viruses compete for target cells, with viral load data from ferrets infected with RSV followed by IAV after 3 d [27]. Our analyses suggested that IAV may reduce the production rate of RSV in addition to RSV reducing the clearance rate of IAV infected cells. In addition, if the two viruses do not compete for target cells, RSV may also reduce the infection rate of IAV. To evaluate how broad the realm of potential mechanisms of viral coinfection with influenza, we then theoretically evaluated different orders and mechanisms of viral interference and cooperation with varying degrees of interaction for IAV-RV and IAV-SARS-CoV-2 coinfections. Qualitatively comparing the model results with weight loss data from animal studies [33, 37] suggested that a reduction in IAV infected cell clearance led to an increased disease phenotype when IAV infection occurred 2-3 d before RV or SARS-CoV-2. In contrast, when IAV was initiated simultaneously with or *<*2 d after RV, a reduction in the rate of clearance of RV infected cells led to reduced disease severity compared to a single IAV infection. Examining how different mechanisms affect viral load and infected cell dynamics during viral coinfection provides important insights into divergent outcomes in addition to generating novel hypotheses regarding why certain virus orders enhance or reduce disease severity.

## Methods

### Data for RSV-IAV coinfection in ferrets

Viral load data were digitized from a study where ferrets monoinfected or coinfected with a long strain of RSV and/or influenza A/Tasmania/2004/2009 (A[H1N1]pdm09; IAV) [27]. Briefly, groups of 4 ferrets were intranasally infected with 3.5 log_10_ 50% tissue culture infectious dose (TCID_50_) of IAV, 5.0 log_10_ plaque-forming units (PFU) of RSV, or IAV followed 3 d later by RSV. Viral RNA copy number per 100 μl of nasal wash was measured daily for 14 d post infection (pi).

### Data for IAV, RV, and SARS-CoV-2 infections in humans

Viral load data were digitized from studies where humans were experimentally or naturally infected with IAV [48], RV [49], or SARS-CoV-2 [50]. For IAV, human volunteers were experimentally infected intranasally with 4.2 log_10_ TCID_50_ of influenza A/Hong Kong/123/77. Nasal washes were collected daily for 7 d and infectious viral titers determined by TCID_50_. The data used herein were from the patient 4 due to this individual having clear viral growth, peak, and decay phases. For RV [49], 14 human volunteers were intranasally infected with 2.4 log_10_ TCID_50_ of RV and was reported as geometric mean. Nasal washes were collected daily for 5 d. For SARS-CoV-2 [50], the data were from naturally infected patients. Viral loads were sampled from throat swabs and measured in RNA copies/ml. All samples were taken approximately 2-4 d after symptoms. We used patient 8 due to clear viral load dynamics and assumed that the infection was initiated 5 d before the onset of symptoms.

### Data for IAV coinfection with RV or SARS-CoV-2 in mice

Weight loss data were digitized from a study where BALB/c mice were intranasally infected with 7.6 *×* 10^6^ TCID_50_ of RV1B and/or 100 TCID_50_ of influenza A/Puerto Rico/8/1934 (PR8) [33]. Coinfections were initiated simultaneously or sequentially at a 2 d interval (IAV-RV or RV-IAV) [33]. Weight loss was measured daily for 14 d.

Weight loss data were digitized from a study where K18-hACE2 mice were intranasally infected with 1 *×* 10^2^ PFU of influenza A/HKx31 (H3N2) and/or 1 *×* 10^4^ PFU of hCoV-2/human/Liverpool/REMRQ0001/2020 (SARS-CoV-2) [37].Coinfections were examined where IAV was given first followed by SARS-CoV-2 after 3 d. Weight loss was measured daily for 10 d.

### Mathematical model of viral monoinfection

To describe the dynamic interactions between epithelial cells and virus during monoinfection, we used the viral kinetic model in Equations (1)-(4) (reviewed in [51, 52]). Briefly, in the model, target cells (*T*) are infected by the virus (*V*) at a rate *βV* per day. Once virus is internalized, the cell undergoes an eclipse phase (*E*), where infected cells do not yet produce virus. The cells then transitions to infectious phase (*I*) at a rate *k* per day. Productively infected cells are cleared at a rate *δ* per day. Virus is produced at rate *p* per cell per day and cleared at rate *c* per day.

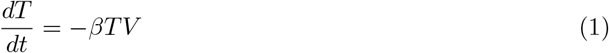

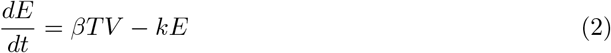

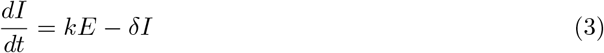

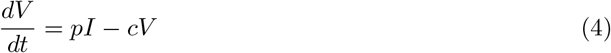

### Mathematical model of viral coinfection

#### Target cell competition

To model viral-viral coinfections, we expanded the model in Equations (1)-(4) using two hypotheses. The first hypothesis assumes that two viruses (*V*_1_ and *V*_2_) compete for target cells (*T*) (‘target cell competition model’; Equations (5)-(8)) [45]. In this model, a single equation for target cells (*T*) was used where each virus can infect these cells at rates *β*_1_*V*_1_ per day and *β*_2_*V*_2_ per day. All other equations are equivalent to those in the monoinfection model. The subscripts *i* = 1, 2 denote each virus.

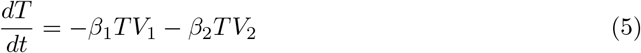

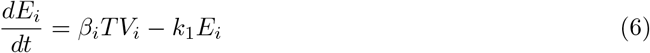

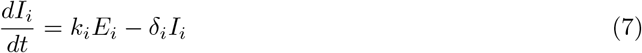

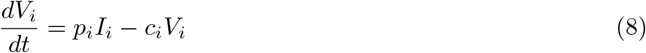

#### Target cell partitioning

The second hypothesis assumes that each virus has its own pool of cells to infect (*T*_1_ and *T*_2_) because viruses may preferentially infect certain cell types [53–57] or be present in a different areas of the respiratory tract (‘target cell partitioning model’; Equations (9)-(12)). All other equations remain the same but are distinct for each virus, resulting a total of 8 equations. The subscripts *i* = 1, 2 denote each virus.

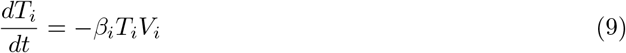

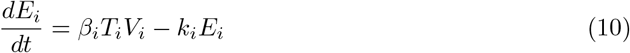

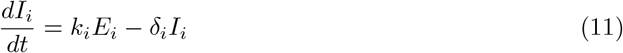

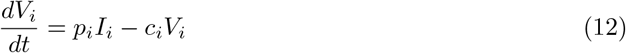

#### Modeling viral-viral interactions

To assess the affect of one virus on another, we used functions that enhance (*α*(*V*_*i*_); Equation (13)) or inhibit (*ζ*(*V*_*i*_); Equation (14)) a particular infection process (i.e., rates of virus infection (*β*_*i*_), virus production (*p*_*i*_), infected cell clearance (*δ*_*i*_), or virus clearance (*c*_*i*_)).

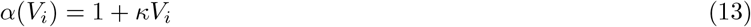

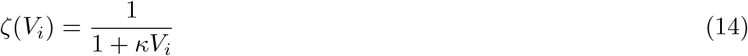

The parameter *κ* is the strength of the interaction.

#### Quantifying the relative change in total virus

To quantify changes in the viral loads and the total viral burden as a consequence of an interaction during coinfection, we calculated the relative change in total virus (i.e., the area under the curve [AUC]) of viral load using Equation (15),

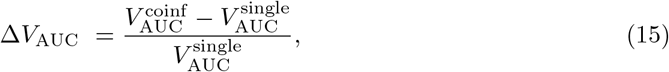

where 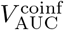 is the AUC for the coinfection and 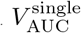 is the AUC for the monoinfection. The AUC was calculated using the Python function *scipy*.*integrate*.*trapz*.

#### Quantifying disease severity

To quantify the percent of the lung infected by the virus, which relates to animal weight loss [46], we calculated the cumulative area under the curve (CAUC) of the infected cell dynamics [46] using the Python function *scipy*.*integrate*.*cumtrapz*.

#### Parameter estimation

For model fits to the ferret data, parameters were estimated using a nonlinear mixed-effect modeling (NLME) and stochastic approximation expectation minimization (SAEM) algorithm implemented in Monolix 2019R1 [58]. For model fits to the human data, parameters were estimated using scipy.optimize.minimize in Python. In the NLME approach, each individual parameter is written as 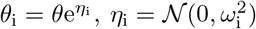, where *θ* denotes the median value of the parameter in the population and *η*_i_ denotes the random effect that accounts for the inter-individual variability of the parameter within the population. Inter-individual variability was allowed for all the estimated parameters with the assumption of no correlation and applying an additive residual error model for log_10_ viral loads.

The initial number of target cells (*T*_0_) was set to 5 *×* 10^7^ cells for ferrets and 2 *×* 10^8^ cells for humans. Similar to our previous studies [46, 59, 60], we fixed the initial number of infected cells (*E*_0_) to 3.1*×*10^3^ cells for IAV infection and 1.0*×*10^5^ cells for RSV infection in ferrets [27] and 1*×*10^2^ cells for all infections in humans. We considered other values of *E*_0_ and found no significant differences in estimated parameters, which is consistent with our prior studies [46, 59, 60]. The initial number of productively infected cells (*I*_0_) and the initial free virus (*V*_0_) were set to 0.

The duration of eclipse phase (1*/k*) for each virus was kept within a biologically feasible value and set to 4.8 h for IAV [59] and to 8.0 h for RSV [61], RV [62], and SARS-CoV-2 [63–65]. For the monoinfection model (Equations (1)-(4)), estimated parameters included the rates of virus infection (*β*), virus production (*p*), virus clearance (*c*), and infected cell clearance (*δ*). The rate of virus infection (*β*) was allowed to vary between 1*×*10^−9^ and 1.0 RNA^−1^ d^−1^ or 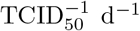, and the rate of virus production (*p*) was allowed to vary between 1*×*10^−3^ and 1*×*10^3^ RNA/cell/d or TCID_50_/cell/d. The rate of infected cell clearance (*δ*) was given a lower limit of 1*×*10^−2^ d^−1^ and an upper limit of 1 *×* 10^3^ d^−1^. For the coinfection models, we first simulated the monoinfection until the day of coinfection then employed the coinfection models (Equations (5)-(8) or Equations (9)-(12)) while incorporating the enhancement or inhibition functions (Equation (13) and/or Equation (14)). The strength of interaction (*κ*) was estimated for each scenario.

Fit quality was assessed using the Akaike Information Criteria with small sample size correction (AIC_c_). The model with the lowest AIC_c_ was considered the best, and ΔAIC_c_ ≤ 2 was considered statistically equivalent [66].

## Results

### Model-predicted mechanisms of RSV-IAV coinfection

We used data from ferrets infected with a long strain of RSV and/or influenza A/Tasmania/2004/2009 (A[H1N1]pdm09; IAV) that had their viral loads measured until 14 d postinfection (pi) [27]. When ferrets were inoculated with RSV followed by IAV 3 d later, morbidity was reduced and IAV titers were slightly lower. To begin examining the interactions between RSV and IAV during coinfection that resulted in these dynamics and establish the baseline parameter values for use in our mathematical models, we first fit the monoinfection model (Equations (1)-(4)) to the data from IAV- or RSV-infected ferrets (Table 1, Figure 2A). This showed a robust fit to each data set and yielded a faster rate of infection (*β*; 1.0 *×* 10^−5^ (RNA/100 μl)^−1^ d^−1^ [RSV] versus 3.3 *×* 10^−6^ (RNA/100 μl)^−1^ d^−1^ [IAV]) and slower rate of virus production (*p*; 3.0 *×* 10^−2^ RNA/100 μl/cell/d [RSV] versus 2.5 *×* 10^1^ RNA/100 μl/cell/d [IAV]) for RSV.

**Table 1.** Best-fit parameters for the monoinfection model. Best-fit parameters obtained from fitting the monoinfection model (Equations (1)-(4)) to viral titers from ferrets intranasally infected with IAV at 3.5 log_10_ TCID_50_ or with RSV at 5.0 log_10_ PFU [27]. Parameters are reported as the population median and the standard deviation of the associated random effect (*ω*). The initial numbers of target cells (*T*_0_) and infected cells (*E*_0_) were fixed to the indicated values, and the initial number of productively infected cells (*I*(0)) and the initial virus amount (*V* (0)) were set to 0.

| Parameter | Description | Units | IAV ($\omega$) | RSV ($\omega$) |
| --- | --- | --- | --- | --- |
| $\beta$ | Virus infectivity | $(\text{RNA/100 } \mu\text{l})^{-1} \text{ d}^{-1}$ | $3.3 \times 10^{-6} \text{ (0.6)}$ | $1.0 \times 10^{-5} \text{ (0.04)}$ |
| $k$ | Eclipse phase | $\text{d}^{-1}$ | 5.0 | 3.0 |
| $\delta$ | Infected cell clearance | $\text{d}^{-1}$ | 0.9 (0.02) | 1.1 (1.8) |
| $p$ | Virus production | $(\text{RNA/100 } \mu\text{l}) \text{ cell}^{-1} \text{ d}^{-1}$ | $2.5 \times 10^1 \text{ (0.2)}$ | $3.0 \times 10^{-2} \text{ (0.03)}$ |
| $c$ | Virus clearance | $\text{d}^{-1}$ | 2.0 (0.06) | 1.7 (0.5) |
| $T_0$ | Initial target cells | cells | $5 \times 10^7$ | $5 \times 10^7$ |
| $E_0$ | Initial eclipse cells | cells | $3.2 \times 10^3$ | $1 \times 10^5$ |
| $I_0$ | Initial infected cells | cells | 0 | 0 |
| $V_0$ | Initial virus | $(\text{RNA/100 } \mu\text{l})$ | 0 | 0 |

Using the single infection parameters, we simulated the ‘target cell competition’ model (Equations (5)-(8)) and the ‘target cell partitioning’ model (Equations (9)-(12))(Figure 1) by first assuming that there were no direct interactions (*κ* = 0). Under this assumption, the target cell partitioning hypothesis performed better than the target cell competition hypothesis (i.e., lower AIC_*c*_; 185.3 [‘partitioning’] versus 189.0 [‘competition’]; Table 2), but the data were not precisely replicated.

**Table 2.**
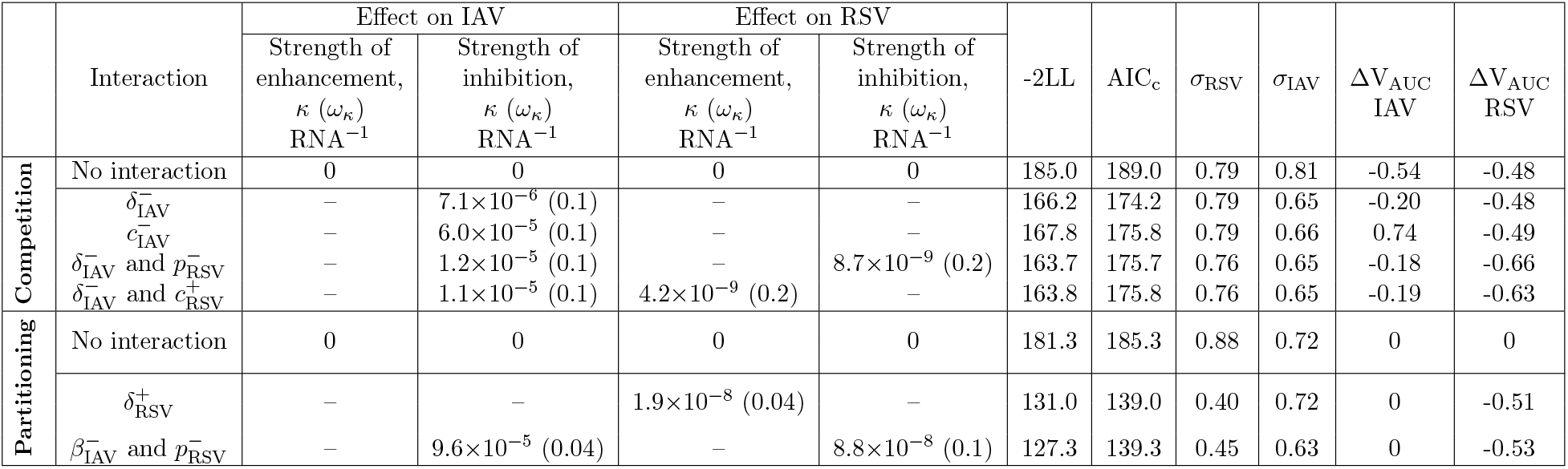
Best-fit parameters of predicted mechanisms of RSV-IAV coinfection. Best-fit parameters from simulating the coinfection models with no interactions (‘no interaction’) or fitting the target cell competition model (‘competition’; Equations (5)-(8)) or the target cell partitioning model (‘partitioning’; Equations (9)-(12)) with the Equation (13) and/or Equation (14) to viral loads from animals infected with RSV followed by IAV after 3 d. An NLME modeling approach was used and only the strength of interaction (*κ*) was estimated. Parameters are reported as the population median (*κ*) with standard deviation of the associated random effect (*ω*_*κ*_). Fit quality is reported as log-likelihood (−2LL), AIC_c_, and standard deviation of the residual error (*σ*). The resulting relative change in total virus (Δ*V*_AUC_) is provided for each scenario.

| | Interaction | Effect on IAV | | Effect on RSV | | -2LL | $\text{AIC}_c$ | $\sigma_{\text{RSV}}$ | $\sigma_{\text{IAV}}$ | $\Delta V_{\text{AUC}}$<br>IAV | $\Delta V_{\text{AUC}}$<br>RSV |
| --- | --- | --- | --- | --- | --- | --- | --- | --- | --- | --- | --- |
| | | Strength of enhancement,<br>$\kappa$ ($\omega_\kappa$)<br>$\text{RNA}^{-1}$ | Strength of inhibition,<br>$\kappa$ ($\omega_\kappa$)<br>$\text{RNA}^{-1}$ | Strength of enhancement,<br>$\kappa$ ($\omega_\kappa$)<br>$\text{RNA}^{-1}$ | Strength of inhibition,<br>$\kappa$ ($\omega_\kappa$)<br>$\text{RNA}^{-1}$ | | | | | | |
| Competition | No interaction | 0 | 0 | 0 | 0 | 185.0 | 189.0 | 0.79 | 0.81 | -0.54 | -0.48 |
| | $\delta_{\text{IAV}}^-$ | – | $7.1 \times 10^{-6}$ (0.1) | – | – | 166.2 | 174.2 | 0.79 | 0.65 | -0.20 | -0.48 |
| | $c_{\text{IAV}}^-$ | – | $6.0 \times 10^{-5}$ (0.1) | – | – | 167.8 | 175.8 | 0.79 | 0.66 | 0.74 | -0.49 |
| | $\delta_{\text{IAV}}^-$ and $p_{\text{RSV}}^-$ | – | $1.2 \times 10^{-5}$ (0.1) | – | $8.7 \times 10^{-9}$ (0.2) | 163.7 | 175.7 | 0.76 | 0.65 | -0.18 | -0.66 |
| | $\delta_{\text{IAV}}^-$ and $c_{\text{RSV}}^+$ | – | $1.1 \times 10^{-5}$ (0.1) | $4.2 \times 10^{-9}$ (0.2) | – | 163.8 | 175.8 | 0.76 | 0.65 | -0.19 | -0.63 |
| Partitioning | No interaction | 0 | 0 | 0 | 0 | 181.3 | 185.3 | 0.88 | 0.72 | 0 | 0 |
| | $\delta_{\text{RSV}}^+$ | – | – | $1.9 \times 10^{-8}$ (0.04) | – | 131.0 | 139.0 | 0.40 | 0.72 | 0 | -0.51 |
| | $\beta_{\text{IAV}}^-$ and $p_{\text{RSV}}^-$ | – | $9.6 \times 10^{-5}$ (0.04) | – | $8.8 \times 10^{-8}$ (0.1) | 127.3 | 139.3 | 0.45 | 0.63 | 0 | -0.53 |

**Figure 1.**
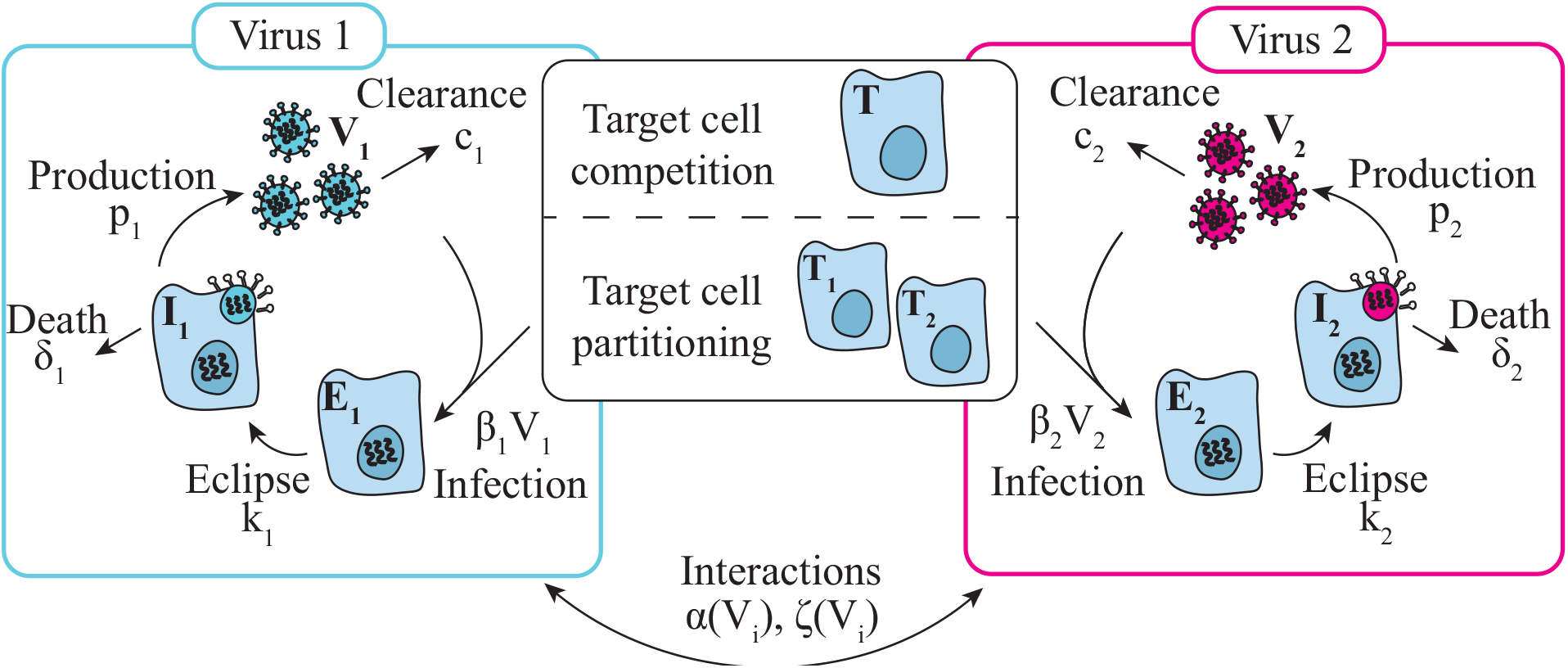
Coinfection model schematic. Schematic of the coinfection models in Equations (5)- and Equations (9)-(12). In the ‘target cell competition’ model, two viruses (*V*_1,2_) interact indirectly by competing for target cells (*T*) [45]. In the ‘target cell partitioning’ model, two viruses do not interact and each have their own pool of target cells (*T*_1,2_). In each model, target cells become infected by virus at rates *β*_*i*_*V*_*i*_, where the subscript *i* = 1, 2 denotes the rates specific to *V*_1_ and *V*_2_, respectively. Infected cells enter an eclipse phase (*E*_*i*_) and transition to producing virus at rate *k*_*i*_. Productively infected cells (*I*_*i*_) produce virus at rate *p*_*i*_ and are cleared at rate *δ*_*i*_. Virus is cleared at rate *c*_*i*_. Direct interactions were implemented by increasing and/or decreasing one or more of the rates using the functions *α*(*V*_*i*_) and/or *ζ*(*V*_*i*_), respectively, due to the other virus.

**Figure 2.**
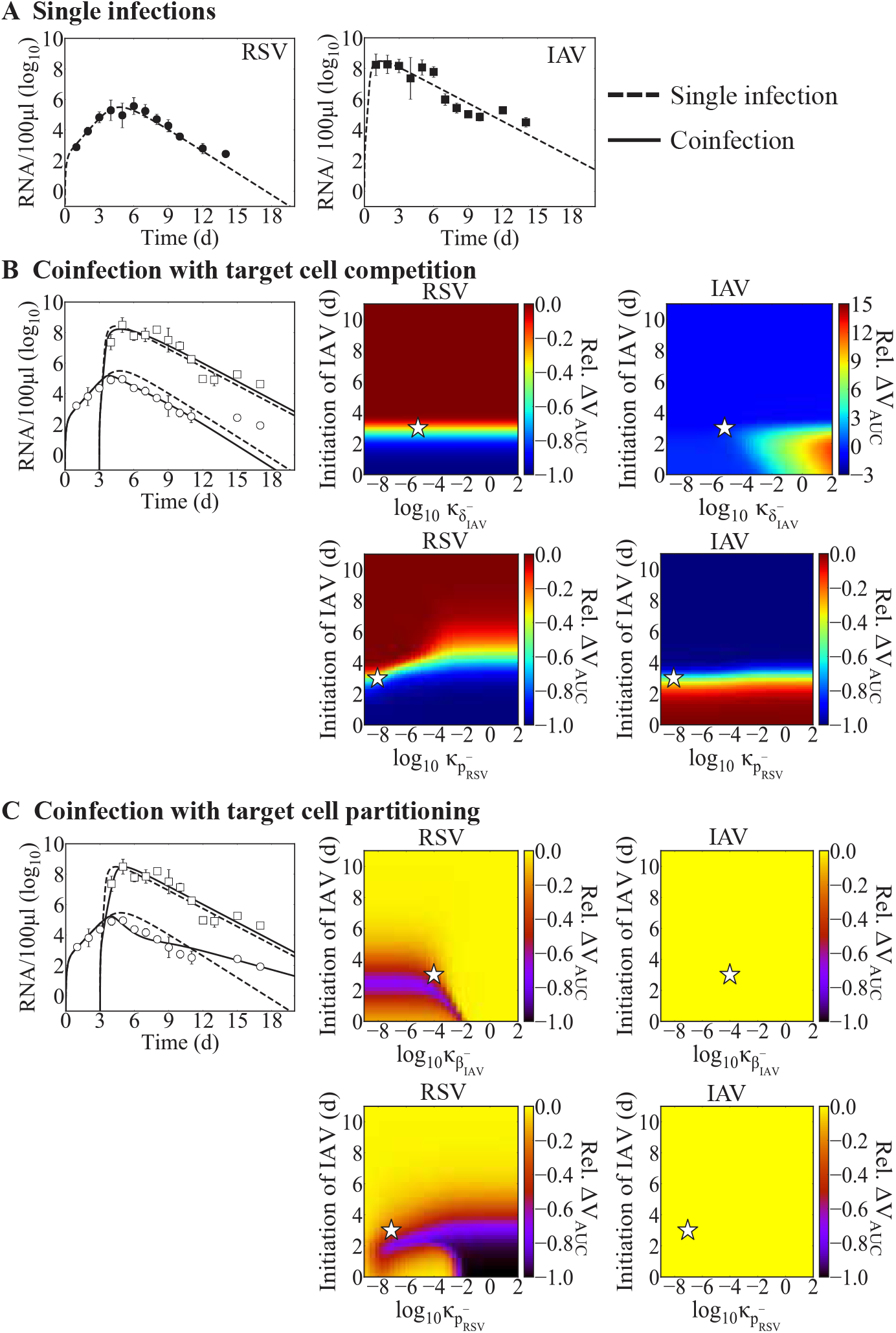
Fit of the RSV-IAV coinfection models. (A) Fit of the single virus model (Equation (1)-(4)) to viral titers from ferrets infected with IAV (black squares) or RSV (black circles). (B-C) Comparison of the single infection model fit (dashed lines) and fit of the coinfection models (solid lines; (B) Equations (5)-(8) or (C)) Equations (9)-(12) with the interaction functions (Equations (13)-(14)) to viral titers from ferrets infected with RSV followed by IAV after 3 d (IAV, white squares; RSV, white circles). Heatmaps are the relative change in total viral burden (i.e., Δ*V*_AUC_; Equation (15)) evaluated for a range of interaction strengths (*κ* = 1 *×*10^−9^ to 1*×* 10^2^ (RNA/100 μl)^−1^) and infection intervals (0 to 11 d). The best-fit *κ* for a coinfection at 3 d is denoted by a white star. Dynamics of the (B) target cell competition model (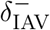 and 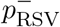) or (C) target cell partitioning model (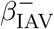 and 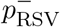).

Thus, to examine whether increases or decreases in the rates of virus infection, production, clearance, or infected cell clearance could better explain the data, we re-fit the models together with Equation (13) and/or Equation (14). In total, we evaluated 16 scenarios for a single interaction and, based on those results, up to 18 scenarios for dual interactions where each virus affected the other (Table S1-S2). When assuming that the two viruses compete for target cells (Equation (5)-(8)), single interactions that resulted in improved fits (i.e., lower AIC_c_) included an RSV-induced reduction in the rate of IAV infected cell clearance 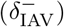 or in the rate of IAV clearance 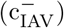 (Table 2; Figure S1A-B). Comparatively, some interactions within the target cell partitioning model (Equations (9)-(12)) led to a rebound of RSV (Figure S2), which were excluded from consideration. The best suggested mechanisms under this hypothesis were either a decrease in the IAV infection rate 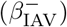 by RSV together with a decrease in the RSV production rate 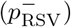 by IAV (Table 2, Figure 1C) or an increase in the rate of RSV infected cell clearance by IAV (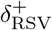; Table S2, Figure S1C).

When allowing for dual interactions, the target cell competition model suggested that there were two sets of mechanisms that provided fits with similar AIC_c_ values as the case when a single interaction was considered (Table 2). Similar to the single interaction results, both sets of mechanisms included a RSV-induced reduction in the rate of IAV infected cell clearance 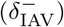. This was paired with either an IAV-induced reduction in the rate of RSV production (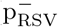; AIC_c_ value of 175.7 (lowest); Figure 2B) or an IAV-induced increase in the rate of RSV clearance (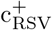; AIC_c_ value of 175.8; Figure S1B). Allowing for dual interactions in the target cell partitioning model suggested an RSV-mediated reduction in the rate of IAV infectivity 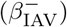 coupled with an IAV-mediated reduction in rate of RSV production (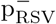; AIC_c_ value of 139.3; Table 2, Figure 2C). The remaining single and double interactions for both models provided fits were not statistically justifiable (Tables S1-S2).

### Effect of infection timing and interaction strength in RSV-IAV coinfection

Because the order, infection interval, and strength of interaction can influence coinfection dynamics and result in diverse disease phenotypes, we sought to better understand how these metrics alter RSV-IAV coinfection. To do this, we evaluated the relative change in total viral burden for each virus (i.e., Δ*V*_AUC_; Equation (15)) across a wide range of infection intervals (0 to 11 d) and interaction strengths (*κ*; 1*×*10^−9^ to 1*×*10^2^ (RNA/100 μl)^−1^) in each model. Here, we focused on the best-fit models with the lowest log-likelihood (Table 2; i.e., 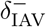 and 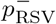 in the target cell competition model and 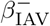 and 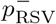 in the target cell partitioning model). For the predicted interactions from the earlier analyses, differing the interaction strength (*κ*) had the strongest effect when the two infections were separated by shorter intervals (i.e., *<*3 d; Figure 2). When the interval was *<*3 d in the target cell competition model, a reduction in the RSV burden (up to Δ*V*_AUC_=-1) was observed along with a prolonged IAV infection (up to Δ*V*_AUC_=15) for an interaction strength higher than the best fit 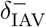 value (white stars in Figure 2B). In the target cell partitioning model, the RSV burden was again reduced but the range of interaction strengths was narrower (Figure 2C). Consistent with the data when the IAV infection was initiated 5 or 7 d after RSV [27], the simulations showed that intervals *>*3 to 4 d after RSV infection resulted in minimal changes in both models (Figure 2B-C). However, only the target cell partitioning led to uninterrupted IAV infection or unchanged IAV viral burden for longer intervals between IAV and RSV infection (Figure 2C).

### IAV coinfection with RV

Animals infected with IAV and RV at the same time or RV two days before IAV yielded lower weight loss and milder disease severity compared to animals infected with IAV alone (Figure 3) [67]. On the contrary, animals infected with IAV 2 d before RV underwent significantly higher weight loss that led to death of all animals by 7 d pi (Figure 3; [67]). To assess the potential mechanisms during IAV coinfection with RV that could lead these empirical observations and provide potential translation to human infection, we first fit the monoinfection model (Equations (1)-(4)) to viral loads from human volunteers infected with IAV [48] or RV [49] (Table 3, Figure 3A). The model fit resulted in different infection kinetic rates for each virus where rate of IAV infection was lower (*β*_IAV_ = 2.5 *×* 10^−6^ (TCID_50_)^−1^ d^−1^ versus *β*_RV_ = 1.6 *×* 10^−3^(TCID_50_)^−1^ d^−1^) and the rate of IAV production was higher (*p*_IAV_ = 3.0 *×* 10^−1^/TCID_50_/cell/d versus *p*_RV_ = 2.9 *×* 10^−3^/TCID_50_/cell/d [RV]).

**Table 3.** Parameters estimates for human infection with IAV, RV, or SARS-CoV-2. Parameter estimates from fitting the single virus model in Equations (1)-(4) to viral load data from humans experimentally infected with 4.2 log_10_ TCID_50_ IAV [68] or 2.4 log_10_ TCID_50_ RV [49], or naturally infected with SARS-CoV-2 [50]. The initial number of target cells (*T*_0_) and infected cells (*E*_0_) were fixed to the indicated values, and the initial number of productively infected cells (*I*_0_) and the initial virus (*V*_0_) were set to 0.

| Parameter | Description | Units | IAV | RV | SARS-CoV-2 |
| --- | --- | --- | --- | --- | --- |
| $\beta$ | Virus infectivity | $[V]^{-1} \text{ d}^{-1}$ | $2.5 \times 10^{-6}$ | $1.6 \times 10^{-3}$ | $1.4 \times 10^{-7}$ |
| $k$ | Eclipse phase | $\text{d}^{-1}$ | 5.0 | 3.0 | 3.0 |
| $\delta$ | Infected cell clearance | $\text{d}^{-1}$ | 3.9 | 5.7 | 4.5 |
| $p$ | Virus production | $[V] \text{ cell}^{-1} \text{ d}^{-1}$ | $3.0 \times 10^{-1}$ | $2.9 \times 10^{-3}$ | 18.8 |
| $c$ | Virus clearance | $\text{d}^{-1}$ | 3.9 | 5.6 | 1.8 |
| $T_0$ | Initial target cells | cells | $2 \times 10^8$ | $2 \times 10^8$ | $2 \times 10^8$ |
| $E_0$ | Initial eclipse cells | cells | 100 | 100 | 100 |
| $I_0$ | Initial infected cells | cells | 0 | 0 | 0 |
| $V_0$ | Initial virus | $[V]$ | 0 | 0 | 0 |
| $\kappa$ | Strength of interaction | $[V]^{-1}$ | See text | See text | See text |

**Figure 3.**
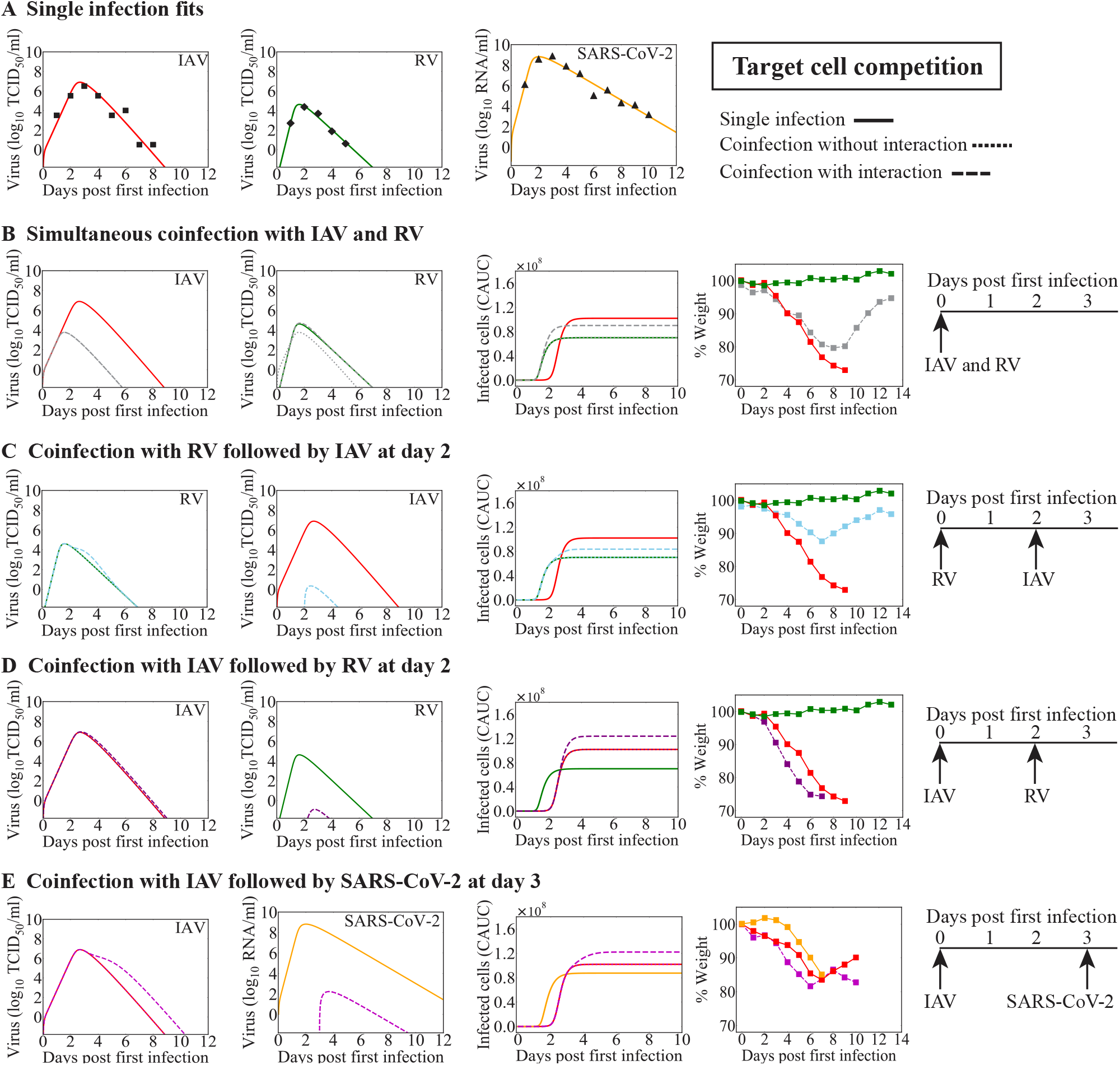
Model predictions coinfection with IAV and RV or SARS-CoV-2 using the target cell competition model. (A) Fit of the monoinfection model (Equation (1)-(4)) to viral titers from humans infected with IAV (red; solid line) [48], RV (green; solid line) [49], or SARS-CoV-2 (yellow; solid line) [50]. (B-E) Model simulations of the dynamics for monoinfection (solid line) or IAV coinfection with RV or SARS-CoV-2 using the target cell competition model (Equations (5)-(8)) without interaction (dotted line) or with interaction (dashed line). The predicted viral loads and CACU of the infected cells are shown alongside the percent weight loss from infected animals [33,37]. (B-C) Dynamics of simultaneous coinfection with IAV and RV or RV-IAV with an IAV-mediated decrease in the rate of RV infected cell clearance 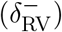. (D) Dynamics of IAV-RV coinfection with an RV-mediated decrease in the rate of IAV infected cell clearance 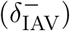. (E) Dynamics of IAV-CoV2 coinfection with a CoV2-mediated decrease in the rate of IAV infected cell clearance 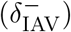.

We next used these parameters in the coinfection models (Equations (5)–(8) or Equations (9)-(12)) with or without interaction (Equation (13) or/and Equation (14)) to predict which mechanisms can lead to the distinct disease outcomes observed in the experimental study. Because viral loads were not measured but weight loss in the infected animals was measured for IAV-RV and RV-IAV coinfections, we compared the estimated cumulative area under the curve (CAUC) of the infected cell dynamics to the weight loss [46], qualitatively matching the magnitude and timing of change. The CAUC of the combined infected cells dynamics of each virus from the coinfection models without any interactions could not recapitulate the weight loss dynamics for any interval or order, confirming that interactions were occurring.

### Simultaneous or sequential RV-IAV coinfection

When testing different interaction mechanisms that enhanced or inhibited one virus within the target cell competition model, the model predicted that the mechanism that could lead to the reduced disease severity observed in RV-IAV coinfection (simultaneous or separated by 2 d) was an IAV-mediated decrease in the rate of RV infected cell clearance (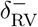; Figure 3B-C). An intermediate signal strength was required for the simultaneous infection (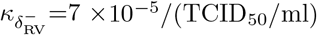; Figure 3B) and a larger signal strength was required for a coinfection separated by 2 d (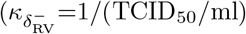; Figure 3C). In both scenarios, this could produce similar reductions in the CAUC of the infected cells as the weight loss patterns in coinfected animals (Figure 3B-C). In addition, when IAV and RV were initiated simultaneously, the model-predicted kinetics showed significant reductions in IAV titers compared to IAV monoinfection (Figure 3B). This was accompanied by a small increase in RV titers around the peak that was due to the IAV-mediated decrease in the rate of RV infected cell clearance. However, in contrast to the simultaneous infection, the model indicated that RV significantly reduced IAV titers when RV was initiated two days before IAV (Figure 3C).

The reduced rate of RV infected cell clearance 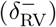 was also identified by the target cell partitioning model (Figure 4A-B). However, for a simultaneous infection, this needed to be coupled with an RV-mediated increase in the rate of IAV infected cell clearance (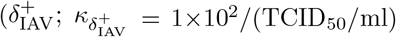; Figure 4A), an increase in the rate of IAV clearance (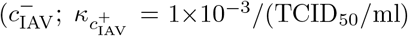; Figure S3A), a decrease in the rate of IAV production (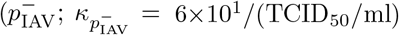; Figure S3B), or a decrease in the rate of IAV infectivity (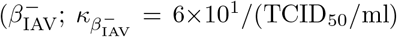; Figure S3C) to achieve a CAUC of the infected cells consistent with the reduced weight loss. In all cases, the combined CAUC of the infected cells was reduced to a level below that of an IAV single infection and accompanied with a complete suppression of IAV titers without affecting RV titers.

**Figure 4.**
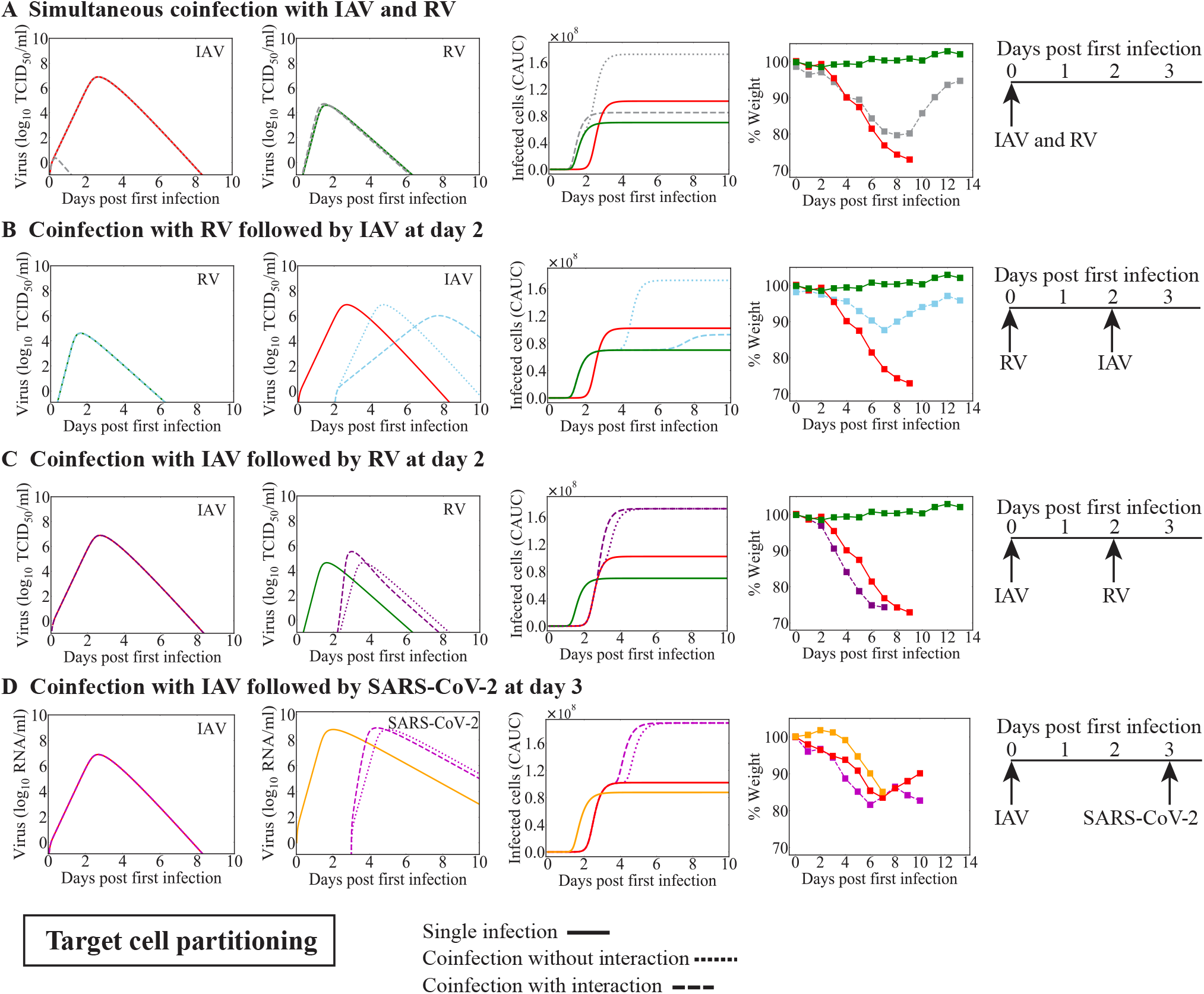
Model predictions for coinfection with IAV and RV or SARS-CoV-2 using the target cell partitioning model. Model simulations of the dynamics for monoinfection (solid line) or IAV coinfection with RV or SARS-CoV-2 using the target cell partitioning model (Equations (9)-(12)) without interaction (dotted line) or with interaction (dashed line). The predicted viral loads and CACU of the infected cells are shown alongside the percent weight loss from infected animals [33, 37]. (A) Dynamics of simultaneous coinfection with IAV and RV with an RV-mediated increase in the rate of IAV infected cell clearance 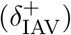 and an IAV-mediated decrease in the rate of RV infected cell clearance 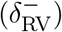. (B) Dynamics of RV-IAV coinfection with reducing the initial number of target cells (*T*_0_) by 1 log_10_ for the second infection. (C) Dynamics of IAV-RV coinfection with an IAV-mediated increase in the rate of RV production 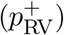. (D) Dynamics of IAV-CoV2 coinfection with IAV-mediated increase in the rate of SARS-CoV-2 production 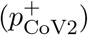.

For RV-IAV coinfection, no single interaction could reproduce the reduced disease severity. Thus, we did not consider dual interactions. However, because virus infections can initiate and/or modify host responses (e.g., type I interferon, macrophages, and neutrophils) that are not included in our model and this can translate into a reduced number of susceptible cells, which is not automatically created by the target cell partitioning hypothesis, we examined the effect indirectly by reducing the initial number of target cells, as done in prior studies [46, 60], that were available for the second virus. For RV-IAV coinfection, reducing the initial number of target cells by 1 log_10_ (i.e., *T*_0_ = 2*×*10^7^ cells for IAV compared to *T*_0_ = 2*×*10^8^ cells for RV; Table 3) was sufficient to reduce the combined CAUC of the infected cells compared to IAV monoinfection (Figures 4B). However, the estimated CAUC of the infected cells deviated from the experimental results at later time points, where the model suggested similar but delayed IAV titers (Figure 4B).

### IAV-RV coinfection

The mechanism that could lead to the increased disease severity observed in IAV-RV coinfection (separated by 2 d) within the target cell competition model was an RV-mediated decrease in the rate of IAV infected cell clearance (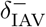; Figure 3D). The required signal strength was large 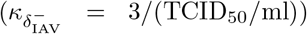. Despite the model-predicted significant reduction in RV titers, the small increase in IAV titers was sufficient to create an increase in the combined CAUC of the infected cells, which aligned with the increased weight loss observed in coinfected animals (Figure 3D).

In contrast, the target cell partitioning hypothesis alone (i.e., no interactions) led to a higher combined CAUC of the infected cells and alignment with the observed increase in disease severity (Figure 4C). This resulted similar viral loads as the monoinfection for both viruses. However, including an IAV-mediated increase in the rate of RV production (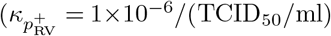; Figure 4C) or the rate of RV infectivity (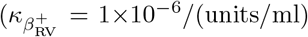; Figure S3D) resulted in an earlier increase in the CAUC of infected cells, which matched the timing of the deviation in weight loss slightly better. In both scenarios, the predicted IAV titer dynamics were similar and unchanged from a monoinfection, and the predicted RV titer dynamics had a similar shape but were much higher when the production rate was increased (5.51 log 10 TCID_50_/ml versus 4.63 log 10 TCID_50_/ml). Other possible mechanisms identified by this model included a reduction in the initial number of target cells (i.e., *T*_0_ = 2 *×* 10^7^ cells for RV compared to 2*×*10^8^ cells for IAV; Table 3) coupled with either a reduction in the rate of IAV infected cell clearance by RV (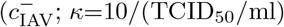; Figure S3E) or in the rate of RV infected cell clearance by IAV (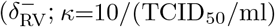; Figure S3F). In the first scenario, the model suggested IAV titers would remain high for an extended period of time, which created an extended, flat viral load peak. In the latter case, the model indicated that there would be no changes to IAV titers and that RV had a slower increase with a lower peak.

### IAV coinfection with SARS-CoV-2

Animals infected with IAV followed 3 d later by SARS-CoV-2 resulted in increased weight loss and more severe disease severity compared to animals infected with IAV or SARS-CoV-2 alone (Figure 3) [37]. To examine the potential interactions between these two viruses, we employed the same approach as above. The results suggested the similar mechanisms for enhanced disease severity for both coinfection models, although the quantitative dynamics were distinct between the models. That is, the target cell competition model predicted a slightly higher combined CAUC of the infected cells when SARS-CoV-2 reduced the rate of IAV infected cell clearance 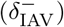 at the signal strength 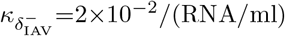 (Figure 3E) where as the target cell partitioning without any interaction led to a significantly higher combined CAUC of the infected cells (Figures 4E). Reducing the initial number of target cells (i.e., a log_10_ reduction [*T*_0_ = 2*×*10^7^ cells]) available to SARS-CoV-2 coupled with a reduction in the rate of SARS-CoV-2-infected cell clearance 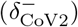 by IAV 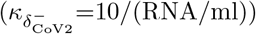 or a reduction in the rate of IAV infected cell clearance 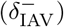 by SARS-CoV-2 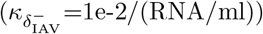 led to a higher combined CAUC of infected cells. The predicted viral load dynamics were distinct between the two models. In the target cell competition model, there was a significant reduction in SARS-CoV-2 titers and a delay in resolution of IAV titers. In contrast, the target cell partitioning model suggested that the SARS-CoV-2 infection was simply delayed.

## Discussion

Respiratory coinfections with multiple viruses are becoming more recognized clinically, particularly in light of the SARS-CoV-2 pandemic. Experimental studies have begun illuminating the outcome heterogeneity, which seems to rely on numerous factors like virus pairing, order and timing of each infection, and specific immune factors. Although the viral and immune dynamics during viral-viral coinfections are only beginning to be defined, mathematical models are useful to predict and narrow the spectrum of potential mechanisms, guide new experiments, and help interpret clinical, experimental, and epidemiological observations. Our analysis on different viral coinfection scenarios suggested that only a small subset of mechanisms could lead to the alterations in viral loads and/or disease severity observed in animal models (summarized in Table 4).

**Table 4.**
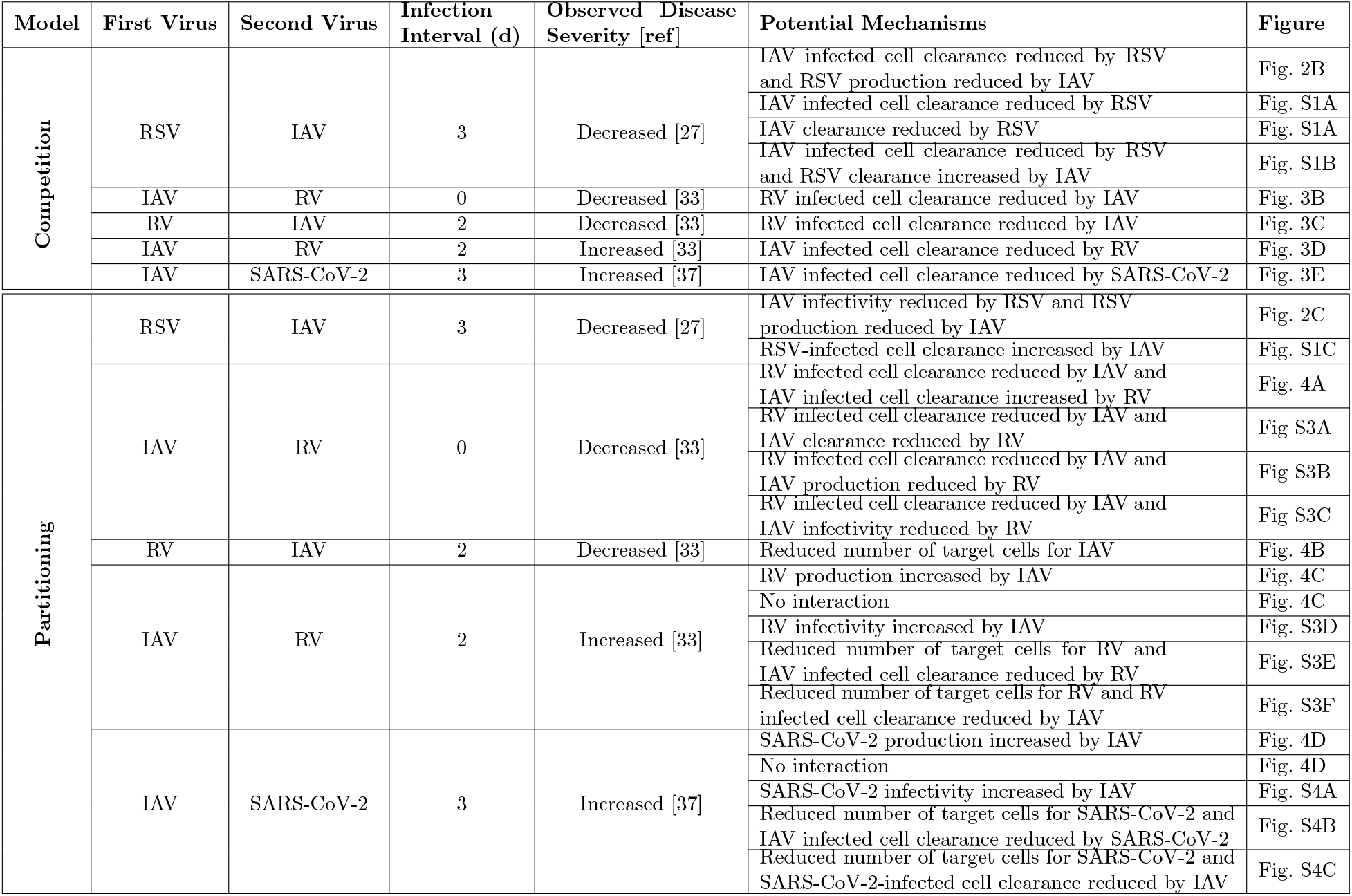
Summary of model-predicted mechanisms resulting in increased or decreased disease severity during viral-viral coinfection. Summary of model-predicted mechanisms that resulted in altered viral loads (RSV-IAV [27]) or disease severity as quantified by the CAUC of the infected cells [46, 47] and measured by animal weight loss (IAV-RV and RV-IAV [33] and IAV-CoV2 [37]).

Early regulation of type I IFNs, macrophages, and/or neutrophils, among other innate immune responses, by a virus could impact the dynamics of subsequent infections. Although we did not assess these immune responses directly, we modeled this indirectly in various ways. The target cell competition model automatically assumes that fewer cells are available for infection by the second virus, which could emulate a protective mechanism that reduces the possible infection size by the coinfecting virus. In the target cell partitioning model, we decreased the number of target cells available for the second virus was reduced, which has been used to mimic lower doses [46, 60]. Both approaches assume that some cells are protected or otherwise unavailable for infection, which can be interpreted as IFN-mediated protection of susceptible cells or an immediate clearance of virus upon infection due to activation of macrophages and/or neutrophils by the first virus. We observed the latter phenomenon in an experiment where CD8 T cells were depleted before infection [46]. In that case, the partial clearance of the inoculum emulated a reduced dose and led to fewer infected cells and, thus, less virus. This automatically reduced inflammation and weight loss [46]. Although we did not examine neutrophils at later time points in that experiment, we would have expected them to be in lower numbers. We did show in separate experiments that neutrophils and macrophages were log-linearly correlated to inflammation [46,47]. Here, allowing for fewer susceptible cells within the target cell partitioning model was sufficient to explain the reduced weight loss in RV-IAV coinfected animals, indicating a similar mechanism. While neither our model nor the data were sophisticated enough to specify the exact mechanism, higher IFN-*β* was detected at 2 d post-coinfection and IAV titers trended slightly lower compared to monoinfection [33]. A follow-up study suggested that the protection of IAV-mediated disease severity by RV was dependent on IFN and that this contributed to the control of neutrophilic inflammation [35]. These data align nicely with our predictions, which may help to connect the underlying reasoning (i.e., fewer cells becoming infected) with downstream consequences (i.e., reduced immune activation and inflammation).

One of the most common mechanisms defined by our analysis was altered rates of infected cell clearance, which may indicate an effect on virus-specific CD8 T cell responses. Variation in the number and composition of epitope-specific T cells following viral coinfection has been observed for other viral pairs (e.g., lymphocytic choriomeningitis (LCMV) and Pichinde (PICV) viruses) [69]. Here, both models predicted modulation of infected cell clearance rates during coinfection with IAV and RV. When these two viruses were given simultaneously, the results suggested that this rate was reduced in all possible mechanisms for RV. Our analysis also predicted that the rate of IAV infected cell clearance was reduced during RSV-IAV coinfection when assuming viruses compete for target cells. While this may indicate negative regulation of the CD8 T cell response either in number or function, it could also indicate that the reduced number of available target cells from RSV pre-infection would have automatically reduced the number of T cells needed to clear the infection. We previously showed that the ratio of infected cells to CD8 T cells drives infection dynamics [46]. Thus, finding increased or reduced quantities of cells without context of other entities (e.g., virus) could result in misinterpretation of the data. Our analysis detected alterations to the rates of virus infectivity or production for some coinfection scenarios, although this was rarely the sole mechanism. Only in IAV-RV and IAV-CoV2 coinfections within the target cell partitioning model were these potential single interactions, which resulted in similar viral load dynamics (Figures 4C-D; Figures S3D and S4A). This is because the type of model used here cannot typically distinguish between the effects of these processes [70]. There is some evidence that infectivity of SARS-CoV-2 is increased by IAV but not RSV within cell cultures [41, 43]. The underlying mechanism driving this remains unknown, but other studies with IAV and PIV have shown enhanced infection rates with PIV increasing cell-to-cell fusion and, thus, spread of IAV [32]. During IAV-CoV2 coinfection in mice, SARS-CoV-2 titers were decreased while IAV titers were increased [42]. The discrepancy between this coinfection resulting in more infected cells but less virus may be due to the results being obtained *in vitro* versus *in vivo* or to another interaction (e.g., IFN suppression of SARS-CoV-2). Our analyses suggested that SARS-CoV-2 viral loads would be reduced within the target cell competition model (Figure S3E-F) or when the number of susceptible cells was reduced within the target cell partitioning model (Figure S4B-C). This may indicate a role for the IAV-activated innate immune response and/or a lower effective dose of SARS-CoV-2.

In this work, we used several data sets with various types of data (i.e., viral loads or weight loss) to predict potential mechanisms during viral coinfection. The type of data used is important because altered viral loads, immune cells, or cytokines do not always directly equate to differences in disease severity. We previously showed this phenomenon, which is due to underlying nonlinearities, while experimentally validating a model of CD8 T cell responses during influenza [46]. In that work, depleting these cells resulted in reduced weight loss, which was counterintuitive given that infection resolution was delayed. However, as mentioned above, the early immune activation led to a predicted lower effective dose (i.e., fewer cells becoming infected) and, thus, reduced disease severity. Similarly in RSV-IAV coinfection, IAV titers were slightly increased yet less weight loss occurred [27]. This could be due to a similar phenomenon, where the higher viral loads later in infection are insignificant with respect to severity. This highlights that while some mechanisms may occur and alter viral loads, they could be distinct from those that yield distinct outcomes. Our results from matching the qualitative differences in weight loss data, which is a measure of disease severity, for IAV coinfection with RV or SARS-CoV-2 may better represent potential mechanisms with measurable consequences. Many of these also led to predicted differences in viral loads. Some information about mechanism may be able to be deduced from the timing of when weight loss begins to deviate from the monoinfection. In several scenarios, this occurred directly after the initiation of the secondary infection, which suggests that the environment created by the first virus has immediate effects. The timing and strength of different mechanisms can influence the ensuing dynamics. Altering these variables during RSV-IAV coinfection agreed with experimental results that the most robust effects occur when RSV was initiated within 0-3 days and that later timings were similar to a monoinfection (Figure 2) [27, 29, 30]. It is likely but unknown whether different mechanisms act only on certain time scales and with varying degrees (i.e., signal strengths, *κ*), but knowledge of this should occur naturally as more data arises and more detailed models are developed.

The mechanisms suggested by the analysis occasionally differed depending on the underlying model hypothesis (i.e., whether viruses compete for epithelial cells) and, in some cases, resulted in different predicted viral load kinetics. Because most respiratory viruses can infect various types of airway epithelial cells and replicate in the upper and lower respiratory tracts, it is conceivable that each virus would have ample cells to infect. However, by chance or due to airway structure, they may enter the same region and interact on a local level. This may lead to cells coinfected with both viruses, which we did not model explicitly. We indirectly modeled the potential effects of coinfected cells by assuming that the rates of infection and/or production could be different. Interestingly, cellular coinfection was detected during simultaneous infection with RSV and HMPV, where coinfected cells were possible but less likely in the presence of IFN [36]. The same may be true during other coinfections with viruses that are sensitive to IFN antiviral responses. To model the impact of coinfected cells and potential heterogeneity in their prevalence, agent-based models may be better suited than those used here.

Using mathematical models to examine data from viral-viral coinfections allowed us to reduce the number of possible underlying mechanisms that could result in altered viral load kinetics and/or disease severity. Although the models were relatively simple and lacked investigation into specific host immune responses, the analysis provided the infrastructure to integrate immunological models of higher complexity once data becomes available. Models for some immune responses during respiratory virus infections are already being developed and validated with experimental data [46,52,71]. Some of the insight from those studies was integrated here and aided our ability to interpret the small amount of experimental data currently available. However, establishing better methods that can predict disease severity (e.g., as in [46]) will be critical. Our ability to assess the contribution and timescales of different mechanisms to infection kinetics and outcome should increase as more temporal viral load, immunologic, and pathologic data become available. In addition, the hypotheses generated should aid experimental design, ultimately leading to a more complete understanding of respiratory virus coinfection.

## Acknowledgments

This work was supported by NIH grants AI125324 and AI139088 (AMS) and P20GM104420 (CR).

## Notes

### Competing Interest Statement

The authors have declared no competing interest.

