## Supplemental Material for "How Interactions During Viral-Viral Coinfection Can Shape Infection Kinetics"

### Complete results from fitting the coinfection models to data from RSV-IAV coinfection in ferrets.

The complete set of best-fit parameters obtained from fitting the coinfection models ('competition', Equations (5)-(8); 'partitioning', Equations (9)-(12)) with the Equations (13) and/or (14) to viral loads from animals infected with RSV followed by IAV after 3 d is in Tables S1-S2. The top models are highlighted in gray and the dynamics are in Figure 2 and Figure S1. Models with a predicted RSV rebound (Figure S2) were excluded.

### Alternate interactions from simulating the target cell partitioning model for IAV coinfection with RV or SARS-CoV-2.

Dynamics of the target cell partitioning model (Equations (9)-(12)) with alternate mechanisms for simultaneous IAV and RV coinfection (Figure S3A-C), IAV-RV coinfection (Figure S3D-F), and IAV-CoV2 coinfection (Figure S4A-C).

**Table S1. Full parameter results from fitting the target cell competition model to RSV-IAV coinfection in ferrets.** Parameters from simulating ('no interaction') or fitting the target cell competition model ('competition'; Equations (5)-(8)) with the Equations (13) and/or (14) to viral loads from animals infected with RSV followed by IAV after 3 d. An NLME modeling approach was used and only the strength of interaction ($\kappa$) was estimated. Parameters are reported as the population median ($\kappa$) with standard deviation of the associated random effect ($\omega_\kappa$). Fit quality is reported as log-likelihood (-2LL), $\mathrm{AIC_c}$, and standard deviation of the residual error ($\sigma$). The best models and predicted mechanisms are highlighted in gray and summarized in Table 1.

| Interaction | IAV | | RSV | | -2LL | $\mathrm{AIC_c}$ | $\sigma_{RSV}$ | $\sigma_{IAV}$ |
|---|---|---|---|---|---|---|---|---|
| | Strength of enhancement, $\kappa$ ($\omega_\kappa$) $\mathrm{RNA}^{-1}$ | Strength of inhibition, $\kappa$ ($\omega_\kappa$) $\mathrm{RNA}^{-1}$ | Strength of enhancement, $\kappa$ ($\omega_\kappa$) $\mathrm{RNA}^{-1}$ | Strength of inhibition, $\kappa$ ($\omega_\kappa$) $\mathrm{RNA}^{-1}$ | | | | |
| No interaction | 0 | 0 | 0 | 0 | 185.0 | 189.0 | 0.79 | 0.81 |
| Single interaction | | | | | | | | |
| $\beta^{+}_{\mathrm{IAV}}$ | $1.9\times10^{-18}$ (6.5) | – | – | – | 185.0 | 193.0 | 0.79 | 0.81 |
| $p^{+}_{\mathrm{IAV}}$ | $3.4\times10^{-12}$ (2.5) | – | – | – | 185.0 | 193.0 | 0.79 | 0.81 |
| $\delta^{+}_{\mathrm{IAV}}$ | $6.7\times10^{-11}$ (8.5) | – | – | – | 185.0 | 193.0 | 0.79 | 0.81 |
| $c^{+}_{\mathrm{IAV}}$ | $2.2\times10^{-10}$ (3.5) | – | – | – | 185.0 | 193.0 | 0.79 | 0.81 |
| $\beta^{-}_{\mathrm{IAV}}$ | – | $4.3\times10^{-18}$ (6.9) | – | – | 185.0 | 193.0 | 0.79 | 0.81 |
| $p^{-}_{\mathrm{IAV}}$ | – | $8.5\times10^{-17}$ (5.3) | – | – | 185.0 | 193.0 | 0.79 | 0.81 |
| $\delta^{-}_{\mathrm{IAV}}$ | – | $7.1\times10^{-6}$ (0.1) | – | – | 166.2 | 174.2 | 0.79 | 0.65 |
| $c^{-}_{\mathrm{IAV}}$ | – | $6.0\times10^{-5}$ (0.1) | – | – | 167.8 | 175.8 | 0.79 | 0.66 |
| $\beta^{+}_{\mathrm{RSV}}$ | – | – | $1.6\times10^{-16}$ (4.5) | – | 185.0 | 193.0 | 0.79 | 0.81 |
| $p^{+}_{\mathrm{RSV}}$ | – | – | $1.2\times10^{-13}$ (2.0) | – | 185.0 | 193.0 | 0.79 | 0.81 |
| $\delta^{+}_{\mathrm{RSV}}$ | – | – | $3.3\times10^{-14}$ (2.0) | – | 185.0 | 193.0 | 0.79 | 0.81 |
| $c^{+}_{\mathrm{RSV}}$ | – | – | $8.6\times10^{-9}$ (0.2) | – | 182.5 | 190.5 | 0.76 | 0.81 |
| $\beta^{-}_{\mathrm{RSV}}$ | – | – | – | $1.4\times10^{-5}$ (2.3) | 182.0 | 190.0 | 0.81 | 0.76 |
| $p^{-}_{\mathrm{RSV}}$ | – | – | – | $1.6\times10^{-8}$ (0.4) | 182.5 | 190.5 | 0.76 | 0.80 |
| $\delta^{-}_{\mathrm{RSV}}$ | – | – | – | $1.1\times10^{-9}$ (0.9) | 183.7 | 191.7 | 0.77 | 0.81 |
| $c^{-}_{\mathrm{RSV}}$ | – | – | – | $2.2\times10^{-13}$ (2.2) | 185.0 | 193.0 | 0.79 | 0.81 |
| Double interactions | | | | | | | | |
| $\delta^{-}_{\mathrm{IAV}}$ and $\beta^{-}_{\mathrm{RSV}}$ | – | $6.5\times10^{-6}$ (0.1) | – | $7.5\times10^{-17}$ (11.1) | 166.1 | 178.1 | 0.79 | 0.65 |
| $\delta^{-}_{\mathrm{IAV}}$ and$p^{-}_{\mathrm{RSV}}$ | – | $1.2\times10^{-5}$ (0.1) | – | $8.7\times10^{-9}$ (0.2) | 163.7 | 175.7 | 0.76 | 0.65 |
| $\delta^{-}_{\mathrm{IAV}}$ and $\delta^{+}_{\mathrm{RSV}}$ | – | $6.6\times10^{-6}$ (0.1) | $3.6\times10^{-8}$ (5.3) | – | 166.1 | 178.1 | 0.79 | 0.65 |
| $\delta^{-}_{\mathrm{IAV}}$ and $c^{+}_{\mathrm{RSV}}$ | – | $1.1\times10^{-5}$ (0.1) | $4.2\times10^{-9}$ (0.2) | – | 163.8 | 175.8 | 0.76 | 0.65 |
| $c^{-}_{\mathrm{IAV}}$ and $\beta^{-}_{\mathrm{RSV}}$ | – | $5.6\times10^{-5}$ (0.1) | – | $8.6\times10^{-21}$ (16.6) | 167.7 | 179.7 | 0.79 | 0.66 |
| $c^{-}_{\mathrm{IAV}}$and $p^{-}_{\mathrm{RSV}}$ | – | $1.0\times10^{-4}$ (0.1) | – | $3.8\times10^{-9}$ (0.3) | 165.5 | 177.5 | 0.76 | 0.66 |
| $c^{-}_{\mathrm{IAV}}$ and $\delta^{+}_{\mathrm{RSV}}$ | – | $6.0\times10^{-5}$ (0.1) | $1.4\times10^{-16}$ (5.1) | – | 167.7 | 179.7 | 0.79 | 0.66 |
| $c^{-}_{\mathrm{IAV}}$ and $c^{+}_{\mathrm{RSV}}$ | – | $9.5\times10^{-5}$ (0.1) | $1.9\times10^{-9}$ (0.3) | – | 165.7 | 177.7 | 0.77 | 0.66 |

**Table S2. Full parameter results from fitting the target cell partitioning model to RSV-IAV coinfection in ferrets.** Parameters from simulating ('no interaction') or fitting the target cell partitioning model ('competition'; Equations (5)-(8)) with the Equations (13) and/or (14) to viral loads from animals infected with RSV followed by IAV after 3 d. An NLME modeling approach was used and only the strength of interaction ($\kappa$) was estimated. Parameters are reported as the population median ($\kappa$) with standard deviation of the associated random effect ($\omega_\kappa$). Fit quality is reported as log-likelihood (-2LL), $AIC_c$, and standard deviation of the residual error ($\sigma$). The best models and predicted mechanisms are highlighted in gray and summarized in Table 1. Although some models resulted in the lower $AIC_c$ values, they were excluded because they produced a viral load rebound.

| Interaction | IAV | | RSV | | -2LL | $AIC_c$ | $\sigma_{RSV}$ | $\sigma_{IAV}$ |
|---|---|---|---|---|---|---|---|---|
| | Strength of enhancement, $\kappa$ ($\omega_\kappa$) $RNA^{-1}$ | Strength of inhibition, $\kappa$ ($\omega_\kappa$) $RNA^{-1}$ | Strength of enhancement, $\kappa$ ($\omega_\kappa$) $RNA^{-1}$ | Strength of inhibition, $\kappa$ ($\omega_\kappa$) $RNA^{-1}$ | | | | |
| No interaction | | | | | | | | |
| No interaction | 0 | 0 | 0 | 0 | 181.3 | 185.3 | 0.88 | 0.72 |
| Single interaction | | | | | | | | |
| $\beta^+_{IAV}$ | $6.0\times10^{-12}$ (2.9) | – | – | – | 181.3 | 189.3 | 0.88 | 0.72 |
| $p^+_{IAV}$ | $3.5\times10^{-11}$ (2.2) | – | – | – | 181.3 | 189.3 | 0.88 | 0.72 |
| $\delta^+_{IAV}$ | $2.3\times10^{-15}$ (2.3) | – | – | – | 181.3 | 189.3 | 0.88 | 0.72 |
| $c^+_{IAV}$ | $7.1\times10^{-16}$ (2.2) | – | – | – | 181.3 | 189.3 | 0.88 | 0.72 |
| $\beta^-_{IAV}$ | – | $1.3\times10^{-6}$ (0.90) | – | – | 180.5 | 188.5 | 0.88 | 0.71 |
| $p^-_{IAV}$ | – | $6.3\times10^{-5}$ (0.06) | – | – | 168.6 | 176.6 | 0.88 | 0.62 |
| $\delta^-_{IAV}$ | – | $7.0\times10^{-7}$ (0.2) | – | – | 178.3 | 186.3 | 0.88 | 0.70 |
| $c^-_{IAV}$ | – | $4.9\times10^{-7}$ (1.9) | – | – | 180.7 | 188.7 | 0.88 | 0.71 |
| $\beta^+_{RSV}$ | – | – | $7.4\times10^{-7}$ (1.0) | – | 178.8 | 186.8 | 0.84 | 0.72 |
| $p^+_{RSV}$ | – | – | $1.4\times10^{-34}$ (12.6) | – | 181.2 | 189.2 | 0.88 | 0.72 |
| $\delta^+_{RSV}$ | – | – | $1.9\times10^{-8}$ (0.04) | – | 131.0 | 139.0 | 0.40 | 0.72 |
| $c^+_{RSV}$ | – | – | $7.1\times10^{-9}$ (0.10) | – | 169.2 | 177.2 | 0.72 | 0.72 |
| $\beta^-_{RSV}$ | – | – | – | $1.8\times10^{-5}$ (0.1) | 128.6 | 136.6 | 0.38 | 0.72 |
| $p^-_{RSV}$ | – | – | – | $1.0\times10^{-8}$ (0.1) | 169.7 | 177.7 | 0.73 | 0.72 |
| $\delta^-_{RSV}$ | – | – | – | $6.8\times10^{-19}$ (4.1) | 181.3 | 189.3 | 0.88 | 0.72 |
| $c^-_{RSV}$ | – | – | – | $8.8\times10^{-18}$ (6.5) | 181.3 | 189.3 | 0.88 | 0.72 |
| Double interactions | | | | | | | | |
| $\delta^-_{IAV}$ and $\beta^-_{RSV}$ | – | $2.1\times10^{-6}$ (0.3) | – | $1.2\times10^{-5}$ (0.1) | 125.9 | 137.9 | 0.38 | 0.70 |
| $\delta^-_{IAV}$ and $p^-_{RSV}$ | – | $2.5\times10^{-6}$ (0.2) | – | $1.0\times10^{-8}$ (0.08) | 164.9 | 176.9 | 0.70 | 0.70 |
| $\delta^-_{IAV}$ and $\delta^+_{RSV}$ | – | $2.8\times10^{-31}$ (10.9) | $1.9\times10^{-8}$ (0.05) | – | 131.0 | 143.0 | 0.40 | 0.72 |
| $\delta^-_{IAV}$ and $c^+_{RSV}$ | – | $2.4\times10^{-6}$ (0.1) | $6.3\times10^{-9}$ (0.1) | – | 164.0 | 176.0 | 0.70 | 0.70 |
| $c^-_{IAV}$ and $\beta^-_{RSV}$ | – | $2.8\times10^{-16}$ (7.2) | – | $1.9\times10^{-5}$ (0.2) | 128.5 | 140.5 | 0.38 | 0.72 |
| $c^-_{IAV}$ and $p^-_{RSV}$ | – | $2.3\times10^{-5}$ (0.3) | – | $6.1\times10^{-9}$ (0.1) | 163.9 | 175.9 | 0.66 | 0.72 |
| $c^-_{IAV}$ and $\delta^+_{RSV}$ | – | $6.0\times10^{-17}$ (3.0) | $1.9\times10^{-8}$ (0.03) | – | 130.9 | 142.9 | 0.40 | 0.72 |
| $c^-_{IAV}$ and $c^+_{RSV}$ | – | $2.6\times10^{-5}$ (0.2) | $3.6\times10^{-9}$ (0.07) | – | 163.1 | 175.1 | 0.65 | 0.73 |
| $\beta^-_{IAV}$ and $\beta^-_{RSV}$ | – | $5.5\times10^{-5}$ (0.4) | – | $1.2\times10^{-5}$ (0.1) | 120.2 | 132.2 | 0.40 | 0.62 |
| $\beta^-_{IAV}$ and $p^-_{RSV}$ | – | $9.6\times10^{-5}$ (0.04) | – | $8.8\times10^{-8}$ (0.1) | 127.3 | 139.3 | 0.45 | 0.63 |

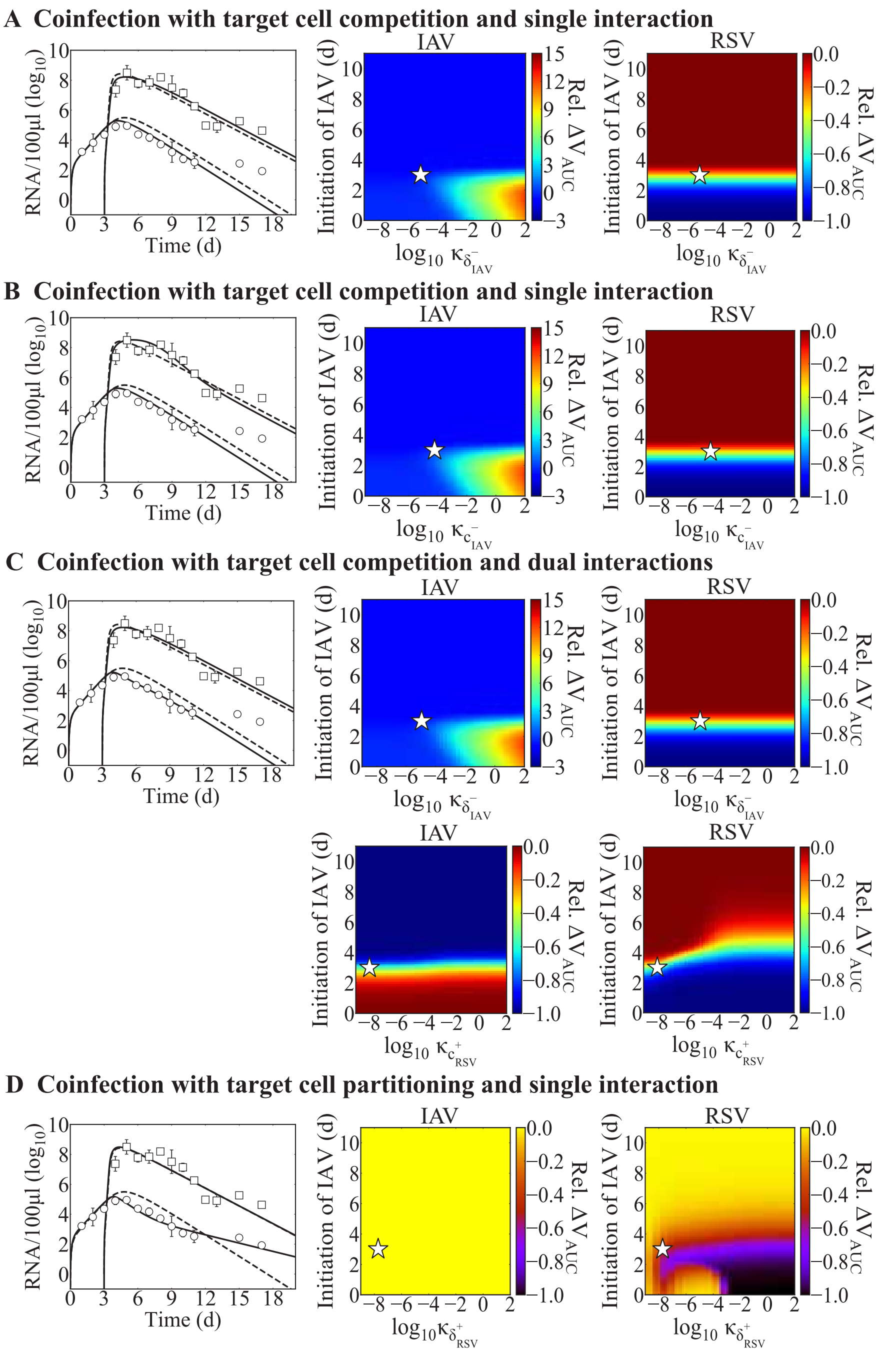


**Figure S1. Model dynamics of alternate mechanisms of RSV-IAV coinfection.** Comparison of the monoinfection model fit (dashed lines; Equations (1)-(4)) and fit of the coinfection models (solid lines; Equations (5)-(8) or Equations (9)-(12)) with the interaction functions (Equations (13)-(14)) to viral titers from ferrets infected with RSV followed by IAV after 3 d (RSV, white circles; IAV, white squares). Heatmaps are the relative change in total viral burden (i.e., $\Delta V_{\text{AUC}}$; Equation (15)) evaluated for a range of interaction strengths ($\kappa = 1{\times}10^{-9}$ to $1{\times}10^{2}$ RNA/100 μl) and infection intervals (0 to 11 d). The best-fit $\kappa$ for a coinfection at 3 d is denoted by a white star. Model dynamics of the target cell competition model with (A) $\delta^-_{\text{IAV}}$ (top) or $c^-_{\text{IAV}}$ (bottom) or (B) $\delta^-_{\text{IAV}}$ and $c^+_{\text{RSV}}$. (C) Model dynamics of the target cell partitioning model with $\delta^+_{RSV}$.

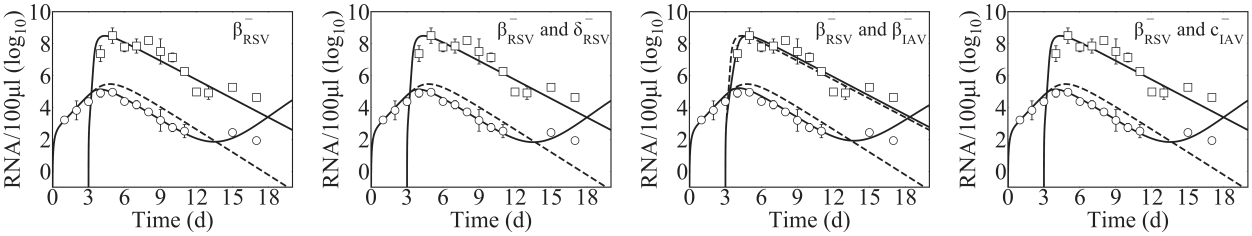


**Figure S2. Model dynamics excluded due to a predicted rebound of RSV during RSV-IAV coinfection.** Fit of the monoinfection model (dashed lines; Equations (1)-(4)) or the target cell partitioning model with the indicated mechanism (solid lines; Equations (9)-(12)) to viral titers from ferrets infected with RSV (circles) followed by IAV (squares) after 3 d. Each mechanism included a reduction in the rate of RSV infectivity ($\beta^{-}_{\text{RSV}}$), which was paired with either decreased rates of RSV infected cell clearance ($\delta^{-}_{\text{RSV}}$), IAV infectivity ($\beta^{-}_{\text{IAV}}$), or IAV clearance ($c^{-}_{\text{IAV}}$). Because the RSV dynamics rebounded, these were excluded as possible mechanisms.

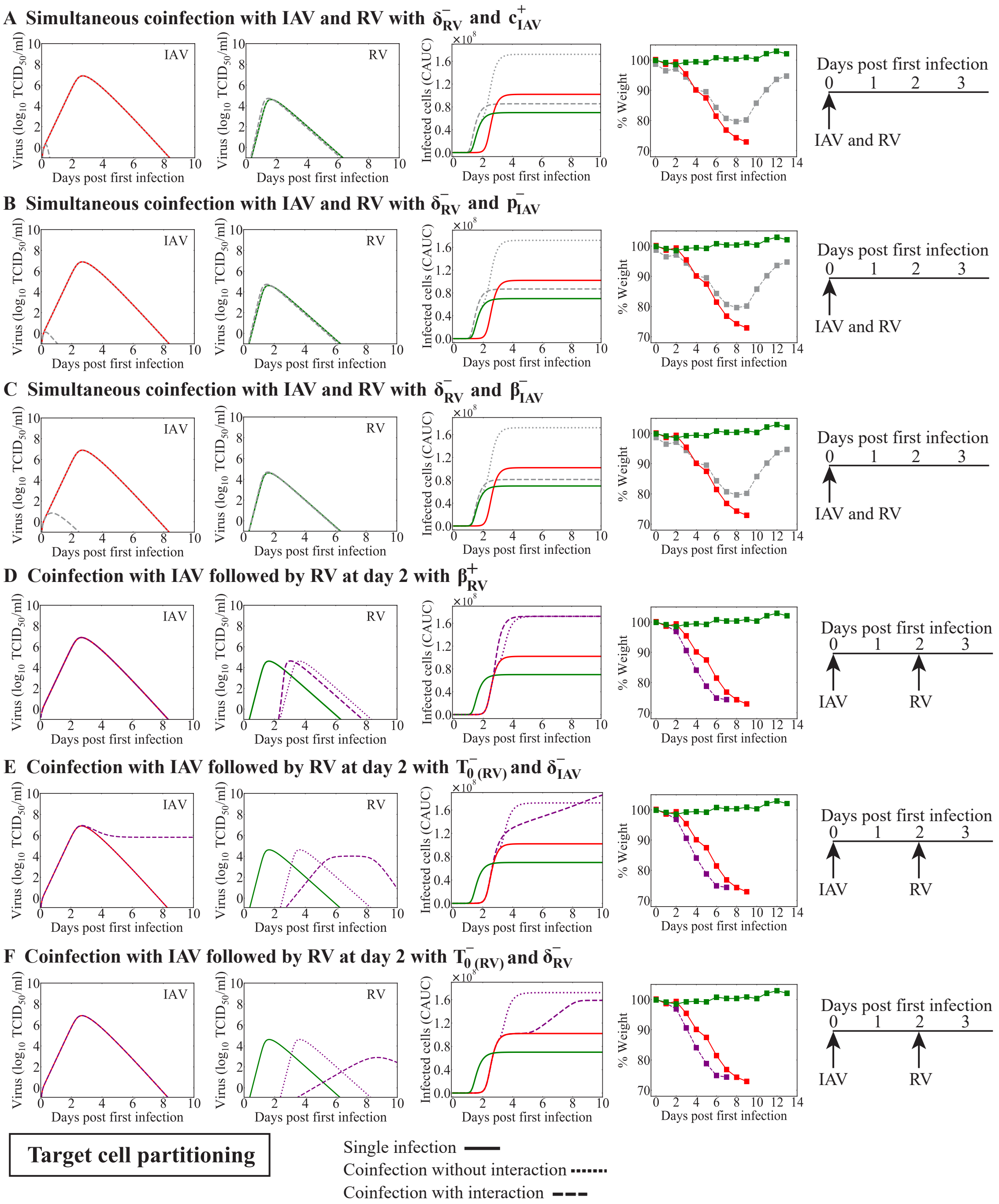


**Figure S3. Model dynamics of alternate mechanisms of IAV-RV coinfection.** Model simulations of the dynamics for monoinfection (solid line) or IAV coinfection with RV using the target cell partitioning model (Equations (9)-(12)) without interaction (dotted line) or with interaction (dashed line). The predicted viral loads and CACU of the infected cells are shown alongside the percent weight loss from infected animals [?]. Dynamics of simultaneous coinfection with IAV and RV with RV infected cell clearance reduced by IAV ($\delta_{\mathrm{RV}}^{-}$) and (A) IAV clearance reduced by RV ($c_{\mathrm{IAV}}^{-}$), (B) IAV production reduced by RV ($p_{\mathrm{IAV}}^{-}$), or (C) IAV infectivity reduced by RV ($\beta_{\mathrm{IAV}}^{-}$). Dynamics of IAV-RV coinfection with (D) RV infectivity increased by IAV ($\beta_{\mathrm{RV}}^{+}$), (E) reduced number of target cells ($T_0^{-}$) for RV and IAV infected cell clearance reduced by RV ($\delta_{\mathrm{IAV}}^{-}$), or (F) reduced number of target cells for RV ($T_0^{-}$) and RV infected cell clearance reduced by IAV ($\delta_{\mathrm{RV}}^{-}$).

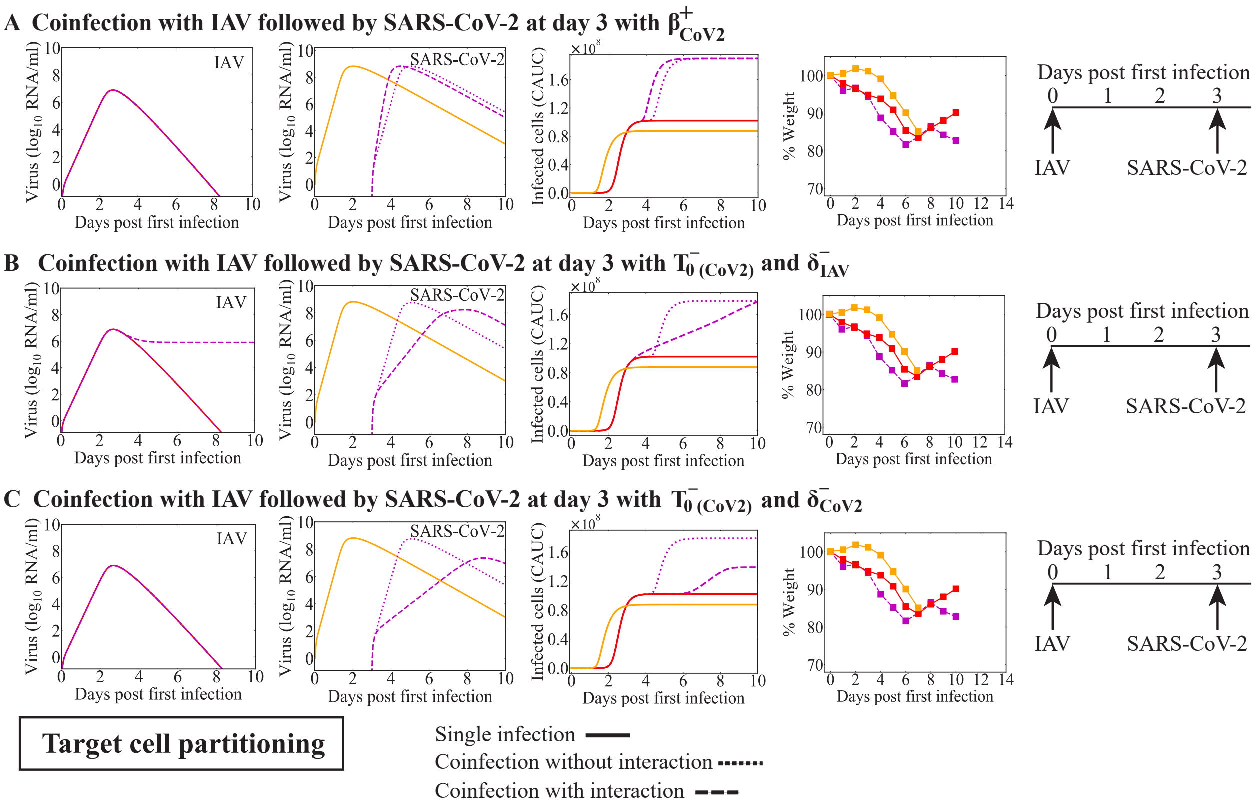


**Figure S4. Model dynamics of alternate mechanisms of IAV-CoV2 coinfection.** Model simulations of the dynamics for monoinfection (solid line) or IAV coinfection with SARS-CoV-2 using the target cell partitioning model (Equations (9)-(12)) without interaction (dotted line) or with interaction (dashed line). The predicted viral loads and CACU of the infected cells are shown alongside the percent weight loss from infected animals [**?**]. Dynamics of IAV-CoV2 coinfection with (A) SARS-CoV-2 infectivity increased by IAV ($\beta^{+}_{\mathrm{CoV2}}$), (B) reduced number of target cells for SARS-CoV-2 ($T_0^{-}$) and IAV infected cell clearance reduced by SARS-CoV-2 ($\delta^{-}_{\mathrm{IAV}}$), or (C) reduced number of target cells for SARS-CoV-2 ($T_0^{-}$) and SARS-CoV-2-infected cell clearance reduced by IAV ($\delta^{-}_{\mathrm{CoV2}}$).
